# Acute exogenous ketone monoester supplementation decreases cardiac vagal modulation in a dose-dependent manner

**DOI:** 10.1101/2025.09.19.677373

**Authors:** Johan S. Thiessen, Aedan J. Rourke, Gaetano C. Pocchi, Claudia M.S. Yong, Addriana Odisho, Jenna A. Nash, Aidan Underwood, Rita Yacoub, Philip J. Millar, Jeremy J. Walsh

## Abstract

Exogenous ketone supplementation can elevate heart rate (HR) but the effects on cardiac autonomic modulation remain unclear. HR variability was analyzed from 18 healthy adults (23 ± 3 yrs; n=10 females) ingesting 0.3 (low-dose) or 0.6 g/kg (high-dose) ketone monoester or placebo in a randomized crossover, double-blind design. Compared to placebo, both ketone doses increased capillary β-hydroxybutyrate and reduced the percentage of successive R-R intervals that differ by ≥50-ms (pRR50) and the root mean square of successive differences (RMSSD) at 45- and 120-min post-ingestion (all p<0.05). Acute ketone ingestion decreases HR variability measures associated with reduced cardiac vagal modulation.

## Introduction

Ketone bodies play an important role in energy metabolism during low carbohydrate intake (Krebs 1966), with more recent work identifying their role as pleiotropic signaling molecules involved in cardiovascular regulation (Lopaschuk & Dyck 2023). Exogenous ketone monoester (KME) supplementation can transiently increase circulating ketone body concentrations without requiring chronic dietary manipulations (Clarke et al. 2012) and has been shown to increase heart rate (HR) (Rourke et al. 2025), stroke volume and cardiac output (Oneglia et al. 2023; Crabtree et al. 2025;), and respiration (McCarthy et al. 2023) in healthy young adults. In contrast, acute KME ingestion does not appear to impact blood pressure (BP) (Marcotte-Chénard et al. 2024).

Despite these intriguing findings, our understanding of how KME ingestion may alter cardiovascular regulation remains poorly described. For example, whether the increases in HR following KME (Oneglia et al. 2023; Crabtree et al. 2025; McCarthy et al. 2021; McCarthy et al. 2023; Rourke et al., 2025) are mediated by underlying changes in cardiac sympathetic or parasympathetic modulation are unclear. Quantifying the variability of R-R intervals in the time- and frequency-domains, otherwise known as HR variability (HRV), can be used to provide insight into underlying modulation of cardiac autonomic activity (Shaffer & Ginsberg 2017). Similarly, indices of BP variability (BPV) are also related to peripheral sympathetic activity (Guerrero et al. 2025). Both HRV and BPV represent non-invasive measures of autonomic modulation and independent predictors of cardiovascular events and end-organ damage (de la Sierra 2023; Shaffer & Ginsberg 2017). Preliminary work reported that KME may decrease time-domain measures of HRV associated with cardiac vagal modulation during hypoxia (Stalmans et al. 2025). However, in this study, KME was ingested before sleep and HRV was measured >10-hours following ingestion. To our knowledge, no study has characterized the time- and dose-dependent effects of acute KME ingestion on HRV or BPV.

Therefore, the purpose of this analysis was to examine the effects of acute KME ingestion on HRV and BPV at rest in young, healthy adults. We hypothesized that acute KME ingestion would decrease HRV without altering BPV in a dose- and time-dependent manner, as compared to placebo. Insights into KME-mediated effects on HRV and BPV will contribute to the growing literature on ketone-mediated cardiovascular regulation.

## Methods

### Ethical Approval and Participants

This study analyzed secondary outcomes from a pre-registered trial (ClinicalTrials.gov NCT06032156) investigating the effects of KME dose on cerebral blood flow (Rourke et al. 2025). The study was approved by the Hamilton Integrated Research Ethics Board (Project ID: 15454) and all participants provided written informed consent prior to commencement. The study recruited individuals 18–35 years of age between September 2023-April 2024 who were not taking ketone supplements or following a ketogenic diet. Those living with obesity (body mass index >30 kg/m^2^) or self-reporting a history of cardiometabolic disease (e.g., diabetes, hypertension), a cardiovascular event requiring hospitalization, or concussion with persistent symptoms were excluded.

### Experimental Protocol

A randomized, counterbalanced cross-over, double-blind experimental design was utilized to assess three conditions: 1) placebo 2) low-KME dose (0.3g /kg^-1^), and 3) high-KME dose (0.6g KME/kg^-1^). The KME was primarily composed of [*R*]-3-hydroxybutyl [*R*]-3-hydroxybutyrate and commercially available (DeltaG Tactical; TdeltaS, Oxford, UK). The placebo solution was prepared using water, denatonium benzoate (NF; Johnson Mathey, London, UK), and xanthan gum to match KME taste and consistency. The placebo solution was added to the KME supplement to ensure each condition was isovolumetric (60 mL). Calorie-free lemon flavouring was used in all conditions to help mask the taste. All supplements were dispensed in white opaque bottles to maintain blinding.

Participants were randomized in a counterbalanced manner by a third-party researcher not involved in data collection. Each visit was separated by ≥72 hours and commenced between 7:00-11:00AM. Participants reported to the laboratory following an overnight fast (excluding water) and abstention from strenuous exercise for 24 hours. The experimental protocol was identical between visits, apart from the consumption of a different supplement (i.e., placebo, low-KME, or high-KME). After arrival at the lab, participants were instrumented with single-lead electrocardiography (lead II; PowerLab Model ML795, ADInstruments, CO, USA) and a non-invasive finger cuff (Human NIBP Nano System, model INL82; AD Instruments, CO, USA) for continuous HR and BP measurements, respectively. Discrete BP was obtained using an automated brachial cuff sphygmomanometer (OMRON Healthcare Co. Ltd, Kyoto, Japan).

All measures were collected in the supine posture following ≥10-minutes of rest. Baseline HR and BP were recorded continuously for 5-minutes. Three discrete brachial cuff measurements were taken during this interval for calibration of continuous BP accuracy. Upon completing baseline measures, participants consumed a supplement drink and rested for 120-minutes, the 0-min timepoint commencing following complete ingestion of the supplement. At 45- and 120-minutes post-ingestion, the collection of HR and BP were repeated. All testing visits were scheduled within ±1 hour to limit diurnal variations.

### Data analysis

Data were analyzed blinded to condition. Indices of HRV was analyzed from 5-minute epochs using the HRV Module in LabChart (ADInstruments, New South Wales, Australia). Time-domain measures included the percentage of successive R-R intervals that differ by ≥50-ms (pRR50), root mean square of successive differences (RMSSD), and standard deviation of the R-R interval (SDRR). Frequency-domain metrics, calculated using fast Fourier transformation, included the high-frequency power (HF; 0.15-0.45 Hz), low-frequency power (LF; 0.04-0.15 Hz) and low-frequency/high-frequency ratio (LF/HF). The Beat Classifier View in LabChart was used to inspect for ectopic beats.

BPV was calculated, as described previously (O’Brien et al. 2023), using the standard deviation (SD) of mean arterial pressure and average-real-variability (ARV). The latter quantifies the absolute average difference between consecutive (beat-to-beat) BP measurements and is calculated as:

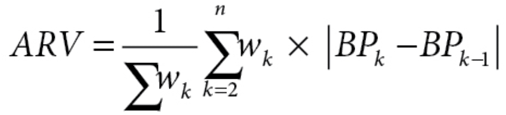

where *n* is the number of BP readings and *w*_*k*_ is the time interval between *BP*_*k*_ and *BP*_*k-1*_.

### Statistical Analysis

A linear mixed-effect model was used to compare measurements between conditions and time (GraphPad Prism v10.1.2, San Diego, CA). Time (baseline vs. 45- and 120-minutes post-ingestion) and condition (placebo, low-KME, high-KME) were included as fixed factors, while participants were included as a random effect to control for within-participant correlations. When appropriate, post-hoc analyses were conducted with a Tukey’s adjustment to control for multiple comparisons. Statistical significance was defined as p<0.05. All data are reported as mean±SD.

## Results

The study recruited 20 participants (10 females; 23±3 yrs, 68.5±10.1 kg, 23.4±2.2 kg/m^2^). Complete HRV data was available for 16 (nine females) participants, with an additional two (one female) participants having complete data from two of the three visits. Complete BPV data was available for 13 participants (seven females) with an additional five (two females) participants having complete data for two of the three visits. In two subjects, the signal quality was unusable for use in HRV and BPV analysis. Ultimately, 18 subjects had complete data for at least two time periods for HRV and BPV analyses.

As reported previously (Rourke et al. 2025), capillary β-hydroxybutyrate concentrations were higher following low and high-KME ingestion compared to the placebo at 45-minutes post-ingestion (3.0±0.8 and 4.1±1.3 vs. 0.1±0.3 mM, both p=0.0001) and 120-minutes post-ingestion (2.4±0.6 and 4.7±0.7 vs. 0.2±0.3 mM, both p<0.0001), but not different from placebo at baseline (0.1±0.1 and 0.1±0.1 vs. 0.2±0.3 mM, both p>0.99). Capillary β-hydroxybutyrate was also higher following high-KME supplementation compared to the low-KME condition at 45- and 120-minute post-supplementation (both p<0.0001). HR increased in a dose-dependent manner by 6±6 and 12±8 beats per minute for the low- and high-dose KME conditions (both p<0.0001, n=18), respectively, while mean arterial pressure remained unchanged across all conditions.

A statistically significant interaction effect was detected for RMSSD, pRR50, and HF power (all p<0.01). At 45-minutes post-ingestion, RMSSD and pRR50 were lower in both low-KME and high-KME compared to placebo. At 120 minutes, they were again lower in both KME conditions compared to placebo, with an additional reduction in high-KME compared to low-KME (Figure 1A and B). Relative HF power also demonstrated an interaction effect (p<0.0001), lower in low-KME and high-KME compared to placebo at 45-min and 120-min post-ingestion. Additional measures of HRV are displayed in Table 1. We did not detect any main effects or interaction effects for relative LF power or the LF/HF ratio (all p>0.09). The beat-to-beat ARV and overall SD of mean arterial pressure showed no interaction or main effects (all p>0.11; Figure 1C and D).

**Table 1.**
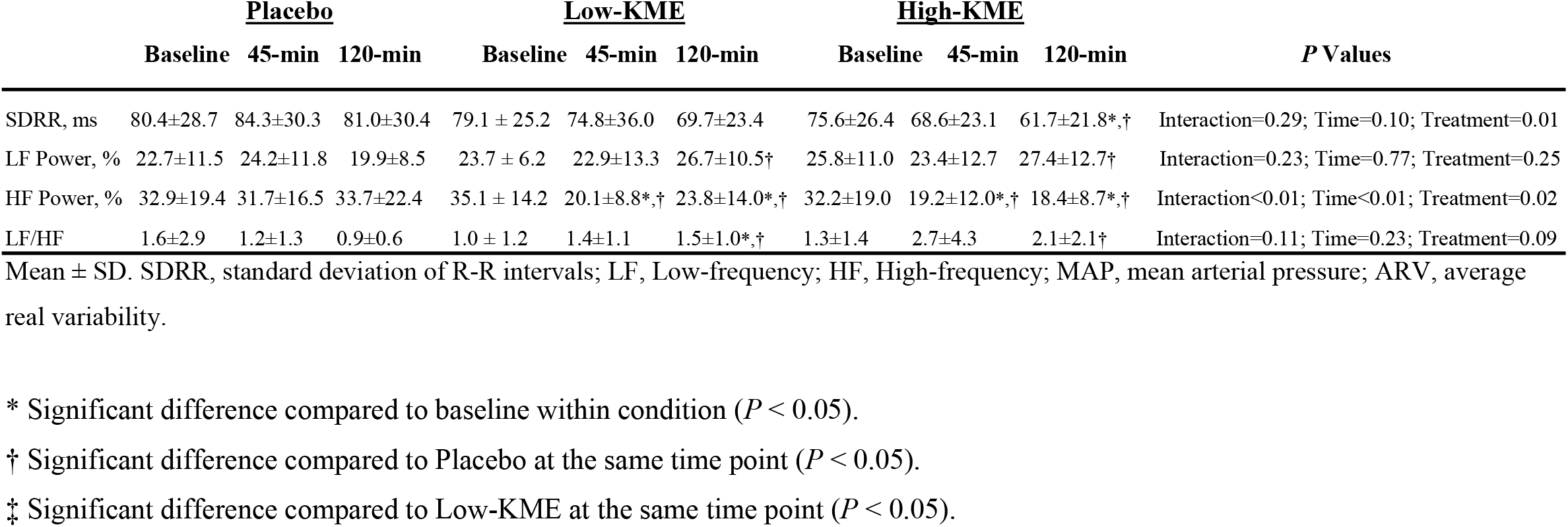
Effects of a low-dose or high-dose acute oral ketone monoester (KME) or placebo ingestion on indices of heart rate and blood pressure variability.

**Figure. 1.**
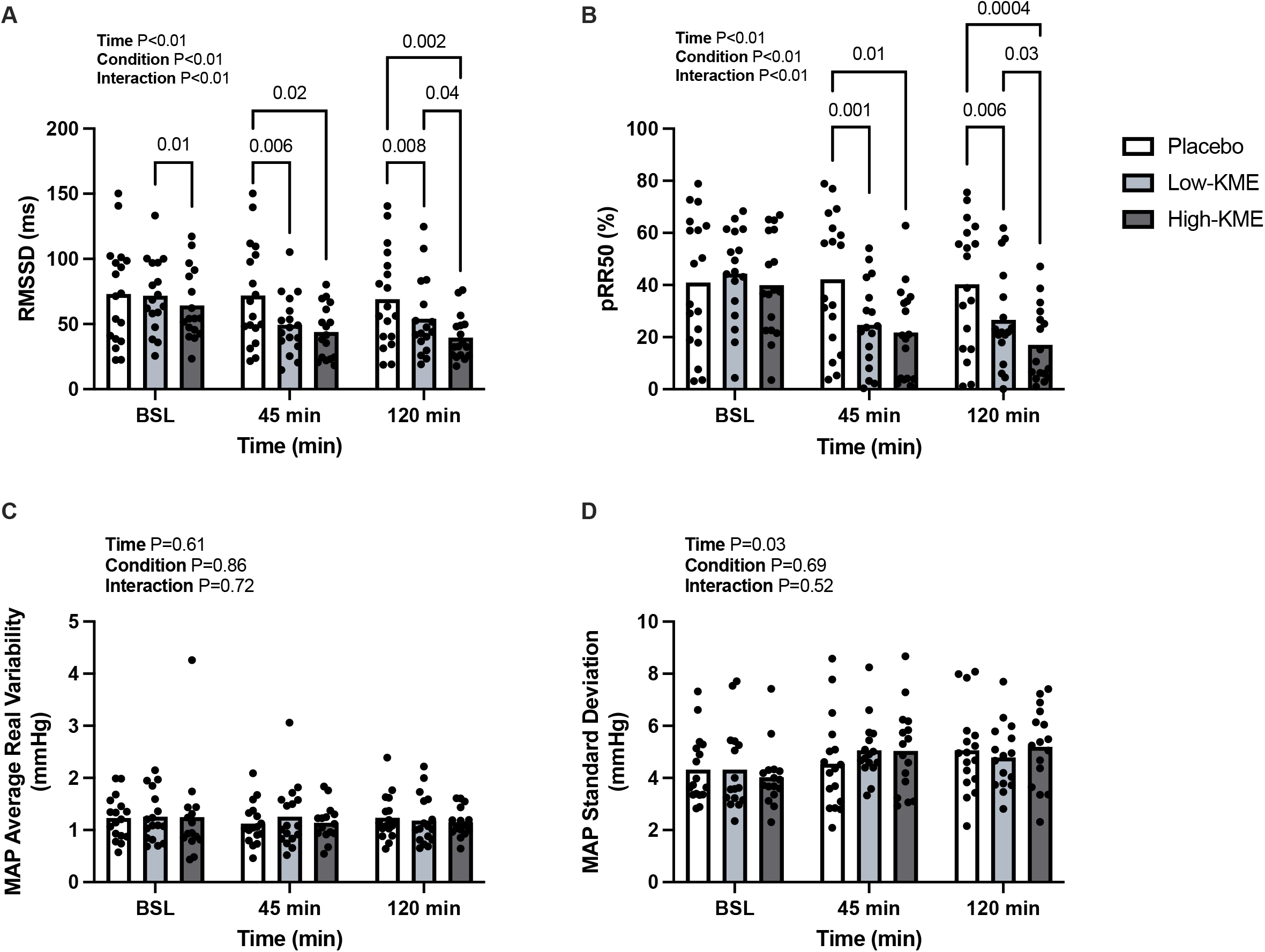

## Discussion

The present analysis examined the dose-dependent effects of acute KME ingestion on HRV and BPV at rest in healthy, young adults. In agreement with our hypothesis, KME ingestion decreased indices of HRV associated with reductions in cardiac vagal modulation (Schaffer & Ginsberg 2017) but did not change BPV. The present data suggests that the hallmark increases in resting HR caused by KME (Oneglia et al. 2023; Crabtree et al. 2025; McCarthy et al. 2021; McCarthy et al. 2023; Rourke et al. 2025) occur secondary to changes in cardiac vagal, not sympathetic modulation. Further, these results demonstrate that BP and its beat-to-beat and overall variability remain unaffected. These results add to a growing body of evidence demonstrating that KME ingestion can influence cardiovascular regulation.

The present findings align with recent work demonstrating that acute 25g KME supplementation 30-minutes prior to sleep at simulated altitude of ∼3,000m lowered HRV indices of cardiac vagal modulation (pRR50 and RMSSD) upon awaking and increased HR (Stalmans et al. 2025). In contrast, intermittent KME supplementation for two days (25g/4-hr during waking hours) decreased HRV during terrestrial hypoxic exposure (∼3,400 m) compared to sea level but not compared to placebo (Narang et al. 2025). The observed differences found between these studies and the present investigation are likely to result from differences in collection timelines (i.e., immediately vs. delayed), dosing strategy (i.e., intermittent vs. acute dosing), and environmental conditions (e.g., hypoxia vs. normoxia).

A hallmark response of KME ingestion is a decrease in blood pH (McCarthy et al. 2021; McCarthy et al. 2023), which may drive chemoreflex-mediated increases in HR (Bruce 1997). Supporting this, we found that KME decreases end-tidal carbon dioxide in a dose-dependent manner (Rourke et al. 2025), consistent with acidosis-induced hyperventilation and chemoreceptor activation (Dempsey et al. 2002). Prior work states chemoreceptor activation increases sympathetic nervous activity (Bruce 1997) and can alter vagal modulation (Somers et al. 1992). KME also increases ventilation which can independently lower vagally-mediated HRV indices (Shaffer & Ginsberg 2017), however, the dose-dependent parallel increases in HR and HRV support changes in vagal modulation, rather than a respiration-only effect. The mechanism in which acute KME ingestion leads to changes in HRV warrants further investigation.

Several methodological considerations should be acknowledged. Our results may not translate to the chronic effects of KME supplementation. Decreased HRV has been reported in those suffering with dietary ketoacidosis (Jaiswal et al. 2012), a condition which chronically increases β-hydroxybutyrate concentrations like those after the high-KME dose (>3 mM). Finally, HRV and BPV are both indirect measures of autonomic control. A recent study reported that acute KME had no impact on muscle sympathetic nerve activity (Seto et al. 2025).

## Conclusion

The present study demonstrates that exogenous ketone supplementation has dose-dependent effects on indices of HRV associated with cardiac vagal modulation, without changing BPV indices or HRV parameters associated with sympathetic activity.

## Author contributions

P.M., J.T., A.R., and J.W. contributed to the conception and design of the work. A.R., C.Y, A.O., J.N., and A.U. contributed to acquisition of the data for the work. J.T., G.P., A.R., R.Y., P.M. and J.W. completed data analysis and interpretation. J.T., P.M., A.R., and J.W. prepared the original manuscript, and G.P. A.R, C.Y, A.O., J.N., A.U., R.Y. and J.W. contributed to revising the manuscript critically for important intellectual content.

## Data Availability

The data that support the findings of this trial are available from the corresponding author upon reasonable request.

## Acknowledgements

The authors would like to thank the dedicated volunteers who participated in this study and the members of the Brain Exercise Enhancement Laboratory who supported data collection. We acknowledge the assistance of Dr. Devin McCarthy and Mr. Elric Alison with early-phase pilot testing for this trial.

